# Reducing the impact of HaHV-1 in Australian abalone: the role of age and immune priming

**DOI:** 10.1101/2025.04.14.648835

**Authors:** Jacinta R Agius, Danielle Ackerly, Angus C Watson, Monique L Smith, Travis Beddoe, Karla J Helbig

## Abstract

Abalone (*Haliotis sp*.) are marine organisms of significant ecological and economic importance. However, disease outbreaks, particularly caused by Haliotid herpesvirus (HaHV-1), pose a major threat to the global aquaculture industry. HaHV-1 causes Abalone Viral Ganglioneuritis (AVG) and has led to significant economic losses due to mass mortality in farmed and wild abalone in regions such as China, Taiwan, and Australia. The current study investigated the effect of age on the susceptibility of Australian hybrid abalone to HaHV-1 and the potential of immune priming as a strategy to protect abalone from HaHV-1 infection. Using a co-housed immersion challenge model, we found that abalone less than one year of age were significantly less susceptible to HaHV-1 infection and exhibit less pronounced clinical signs of HaHV-1 infection when compared to adults. Additionally, immune priming adult abalone with poly(I:C) prior to viral challenge provided protection against HaHV-1 when compared to abalone primed with a bacterial antigen, Flagellin-A and unprimed controls. We also determined that the use of pedal swabs is a less invasive method for confirming positive HaHV-1 infections, but not for determining comparative viral loads. These findings are pivotal in developing preventative strategies against HaHV-1 in aquaculture and highlight the need for further research on immune priming and age-related susceptibility in abalone.

## Introduction

Abalone (*Haliotis sp.*) are marine organisms of ecological and economic importance worldwide. Over the past 20 years there has been a steady increase in the global production of farmed abalone however, one of the major threats to further industry growth is disease outbreaks which are currently estimated to cost the global aquaculture industry US$6 billion annually ^1–3^. Haliotid herpesvirus (HaHV-1) is a large, enveloped DNA virus belonging to the family *Malacoherpesviridae,* which causes the neuropathological disease known as Abalone Viral Ganglioneuritis (AVG) ^4–6^. Outbreaks of AVG have caused significant economic losses due to mass mortality in farmed abalone in China, Taiwan and Australia ^6–10^. HaHV-1 was first detected in Australia on the coast of Victoria in 2005 and has since become recurrent in Victoria, Tasmania and more recently in South Australia ^7,11,12^. Unfortunately, there are no treatments or preventatives currently available against this virus and biosecurity measures are not always feasible, as environmental factors are difficult to control due to the nature of abalone farming systems ^13^.

Following the initial 2005 outbreak and to safeguard Australia’s abalone industry, several studies were conducted to understand HaHV-1 infection in Australian abalone. Most of the published susceptibility data however, has generally been inferred from what was observed during the Victorian outbreak of 2005 ^7^. Although these findings have been imperative in informing the industry about the virus and its associated disease, additional controlled studies are required to make definitive conclusions to appropriately prepare for any future outbreaks. It is currently understood that all Australian abalone tested to date are highly susceptible to HaHV-1 regardless of age and species ^14,15^. Reported clinical signs of infected abalone generally include, irregular peripheral concave elevation of the foot otherwise known as foot curl, excess mucous production, poor adhesion to the substrate, and swollen and protruding mouth parts ^15–18^. Additionally it is important to note that of the limited studies currently available, investigations have generally been conducted on farmed adult hybrid abalone (*H. rubra x H. laevigata)* despite the range of ages (larvae, juveniles, adults) and species (*H. rubra, H. laevigata* & hybrids) present on farm ^19–21^. This necessitates the need for further research to understand how AVG presents in a range of farmed abalone to make informed decisions during outbreaks.

Current means of preventing the spread of AVG are limited to biosecurity methods, for example movement restrictions, quarantine and destruction of diseased stock ^17,18^. These methods are generally actioned once clinical signs are evident, and is too late to restrict pathogen spread in abalone farming systems due to the continuous pumping of seawater into, throughout and out of the farm ^18,22^. This necessitates the need for an alternate strategy against HaHV-1 for farmed abalone. Abalone are invertebrate organisms so therefore lack an adaptive immune response and classical immunological memory which in vertebrates, is afforded by antibodies ^23^. In recent years, several studies have begun to explore the potential of innate immune priming against pathogens in invertebrates. Immune priming is when organisms that do not possess an adaptive immune system can mount an enhanced immune response upon secondary exposure to a pathogen or antigen ^24,25^. This phenomenon has been evidenced in several marine organisms, most notably the penaeid shrimp and Pacific oyster (*Crassostrea gigas*) against white spot syndrome virus (WSSV) and Ostreid herpesvirus (OsHV-1), respectively ^26–30^. In *C. gigas*, injection of an immune stimulant consisting of non-specific and non-infectious dsRNA (poly(I:C)) triggered an anti-viral immune response which as a result protected primed animals against OsHV-1 infection ^28,30,31^. Immune priming is a promising strategy against viruses in economically important marine organisms but has not yet been explored in abalone against HaHV-1.

In this study we utilised a co-housed immersion challenge model to understand the effect of age and immune priming with either poly(I:C) or Flagellin-A protein on HaHV-1 susceptibility in farmed Australian hybrid abalone. To imitate a natural HaHV-1 outbreak on farm, all HaHV-1 challenge trials were conducted with groups of co-housed abalone, as opposed to previous studies where abalone were housed separately following viral challenge. We show that abalone less than one year of age are significantly less susceptible to HaHV-1 infection and that immune priming adult abalone with poly(I:C) prior to subsequent viral challenge, protects against viral infection. The findings of this project are pivotal in the search for a successful preventative strategy against HaHV-1 in aquacultural environments and can provide a starting point for further research against devastating marine pathogens.

## Results

### Investigating the effect of age on HaHV-1 susceptibility

#### Juvenile abalone are less susceptible to HaHV-1 than adult abalone

The ability of abalone age to alter the outcome of HaHV-1 susceptibility has not previously been examined. To begin to understand whether age effects the outcome of HaHV-1 infection the survival of hybrid abalone ranging from 7 to 24 months was measured after challenge with HaHV-1 via immersion. The highest survival rate was observed in 7-month old abalone (∼12%), which differed significantly from 1-year old abalone, having a 50% survival rate (Fig 2A). In comparison the older abalone age groups had significantly increased death rates, with the 1.5-year-olds displaying a 99% death rate at the end of the trial, and the 2-year-old abalone all dying by day 11 (Fig 2A). As shown in Figure 2B and despite the observed difference in survival between age groups, there was no difference in the amount of virus per mg of nerve tissue at death. The general pattern of virus within the nerves of abalone surviving viral challenge indicates that at the point of sacrifice, only 50% of 7-month-olds had detectable levels of virus within their nerves in comparison to ∼83% of the 1-year-old abalone (Fig. 2C). To further investigate if the above trends in survival were only specific to hybrid abalone, the survival of 7-month-old greenlip abalone was measured after HaHV-1 challenge via immersion (Fig. 2D). 7-month-old greenlip abalone were significantly less susceptible to HaHV-1 than 1.5-year old hybrid abalone which reached 0% survival by day 14 as evidenced by the survival trend in Figure 2D.

**Figure 1.**
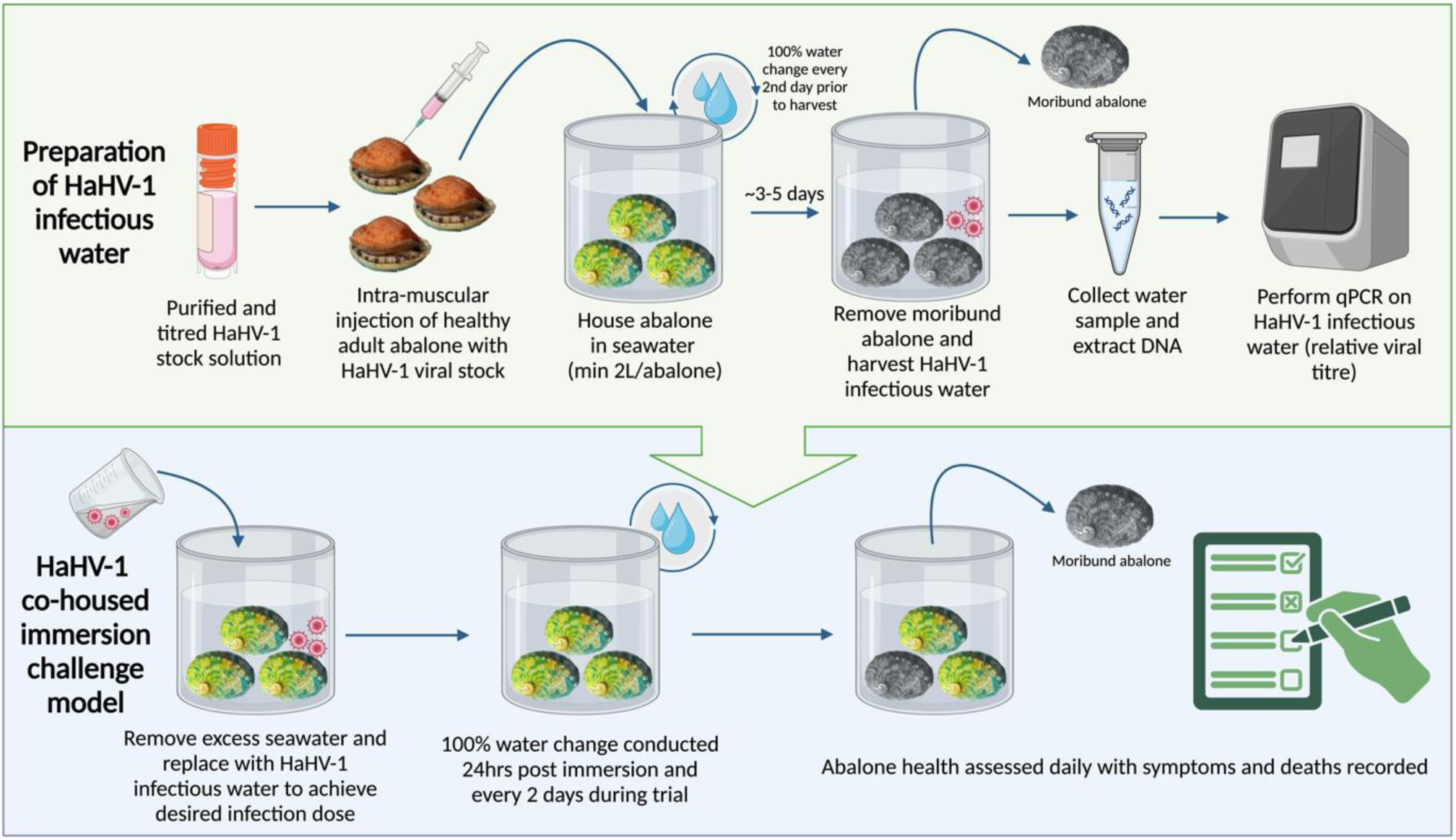
Production of HaHV-1 infectious water for immersion challenge. To produce HaHV-1 infectious water, abalone were first intra-muscularly injected with a purified and titred HaHV-1 stock. Injected abalone were then housed in aquaria (2L seawater/abalone) maintained with 100% water changes. Once abalone were moribund (unable to adhere to the substrate and producing excess mucous), a water sample was taken for subsequent DNA extraction and qPCR analysis to obtain a viral titre. To challenge cohoused groups of naive abalone via immersion, excess seawater was removed and replaced with titred HaHV-1 infectious water to achieve a desired infection dose. 24hrs after immersion, a 100% water change was conducted and again every 2 days thereafter. Abalone health was assessed daily with symptoms and deaths recorded. Created in https://BioRender.com.

**Figure 2.**
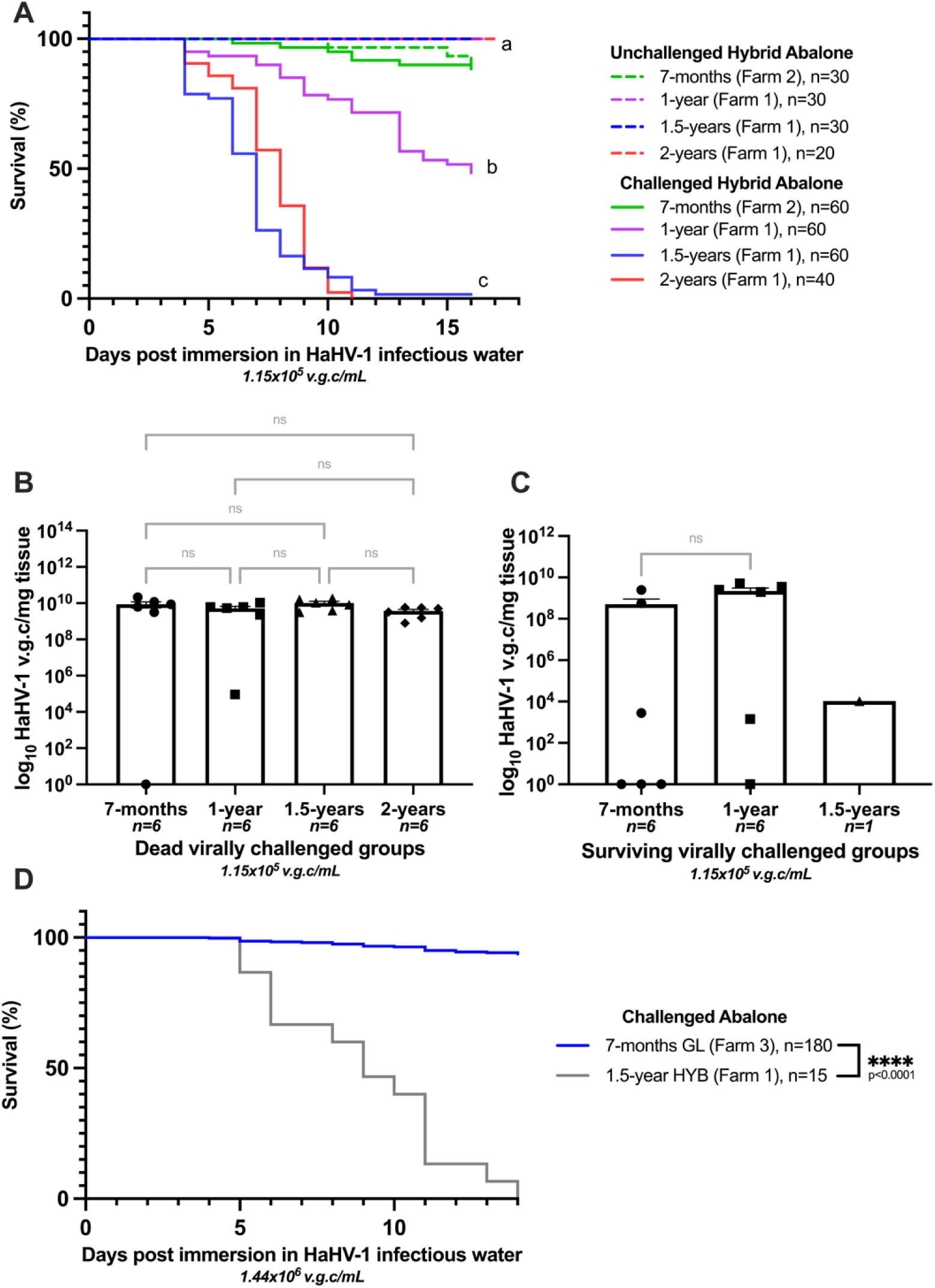
Abalone age impacts susceptibility to HaHV-1. **(a)** Survival of unchallenged (dotted lines) and HaHV-1 challenged (solid lines) abalone aged 7-months to 2-years. Abalone were infected with ∼1.15 x 10^5^ v.g.c/mL via immersion challenge. All experimental abalone groups within an assigned letter show no significant difference between each other, however statistical differences were observed between assigned letters (p<0.0001; Log-rank test, n=20-60). **(b)** Relative viral titre of HaHV-1 per mg of nervous tissue of abalone that died during the trial. Data is represented as mean ± SEM, n=6 abalone, ordinary one-way ANOVA, ns= not significant (p>0.05). **(c)** Relative viral titre of HaHV-1 per mg of nervous tissue of surviving abalone sacrificed at the end of the trial. **(d)** Survival of abalone aged 7-months and 1.5-years after immersion challenge with 1.44 x 10^6^ v.g.c/mL. Log-rank test, n=15-180.

#### Visual symptoms of HaHV-1 differ between juvenile and adult abalone

Older abalone infected with HaHV-1 are known to display symptoms including poor adhesion to the substrate, foot curl, excess mucous production and swollen mouthparts ^15–18^ however, the effect of abalone age on clinical signs of HaHV-1 infection has not been explored. To investigate this, symptoms following exposure to HaHV-1 in abalone 7 to 24 months of age were recorded. In comparison to healthy control abalone, visible symptoms in abalone over 1-year old were more pronounced and reflected previously described clinical signs including foot curl, exposed shell margins and occasionally, prolapse of the mouth ^7,17,18^ (Fig. 3). Conversely, visible symptoms presented after HaHV-1 infection in abalone 7-12 months of age appeared more subtle. As shown in Figure 3, the characteristic HaHV-1 symptom of foot curl was less prominent within this age range, with the most obvious sign of infection being exposed shell margins. Additionally, within experimental housing conditions severe foaming on the surface of the water of abalone over 2.5 years of age is indicative of excessive mucous production and therefore generally an indication of HaHV-1 infection. Interestingly, foaming in the aquaria of infected abalone 1.5-years of age and under was absent (Table 2).

**Figure 3.**
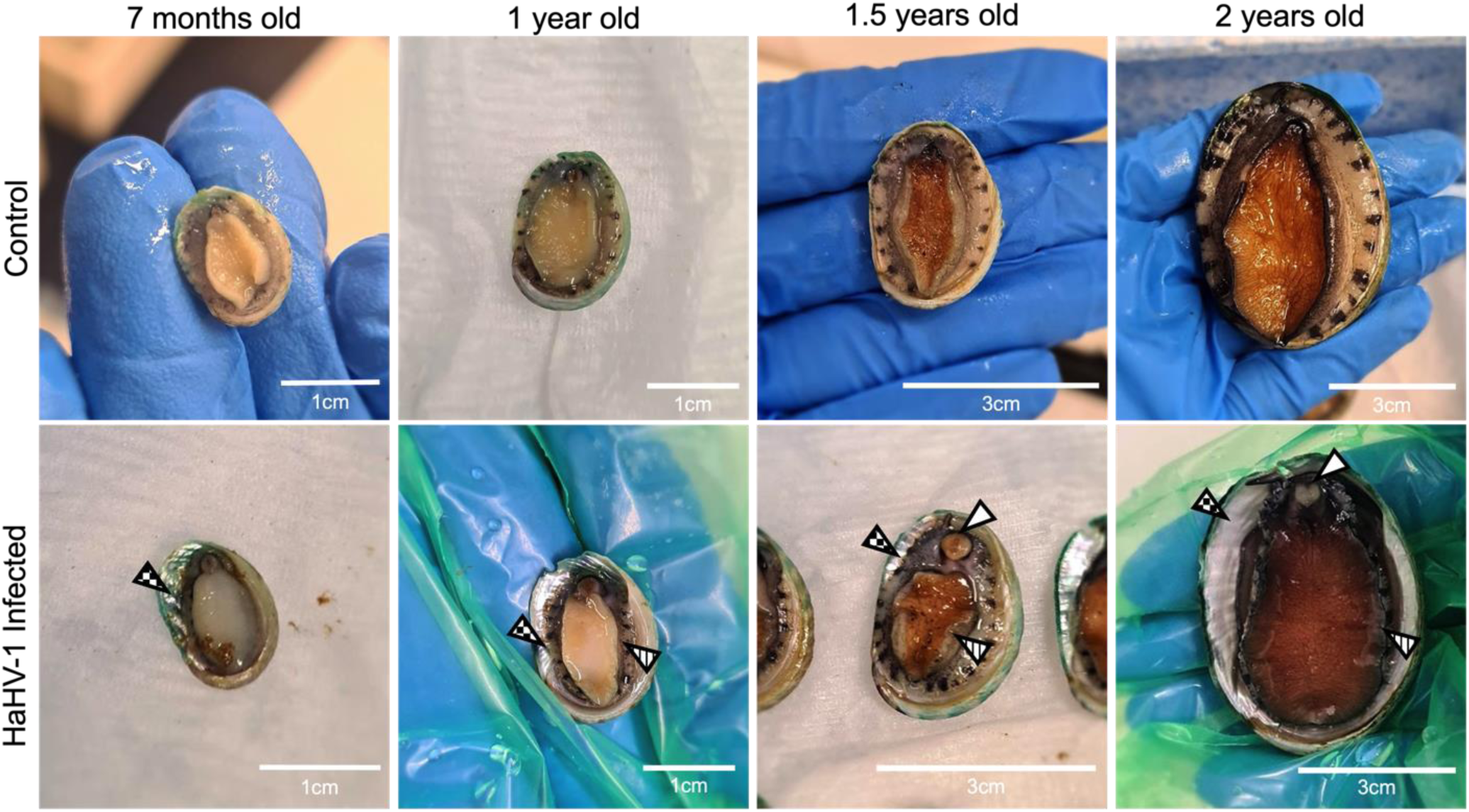
Visible symptoms of HaHV-1 are less pronounced in younger abalone. Images of control and HaHV-1 infected abalone aged 7-months to 2-years. Checkered arrow = exposed shell margins, Vertical striped arrow = foot curl, white arrow = prolapse of the mouth.

**Table 1.**
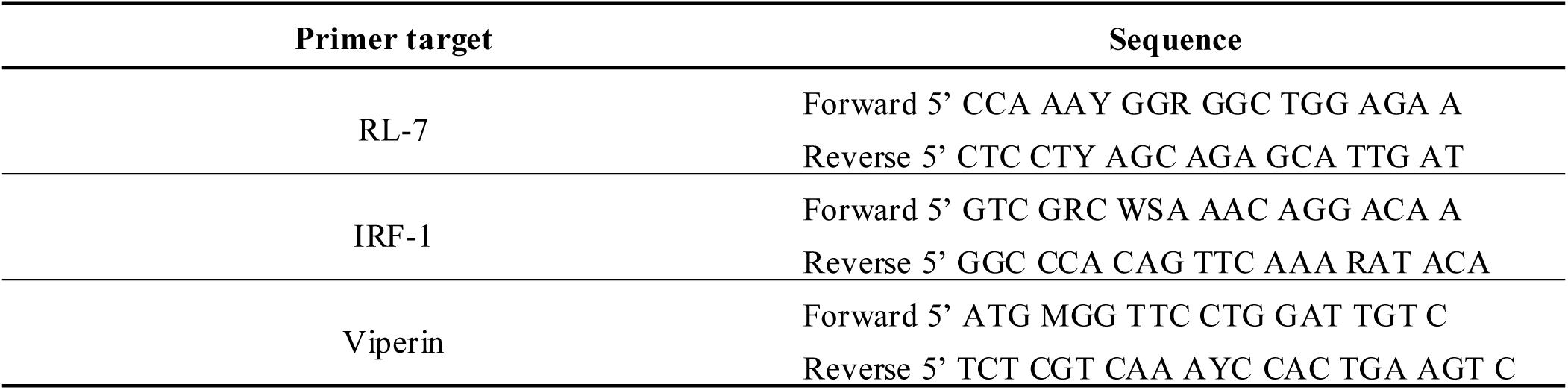

**Table 2.**
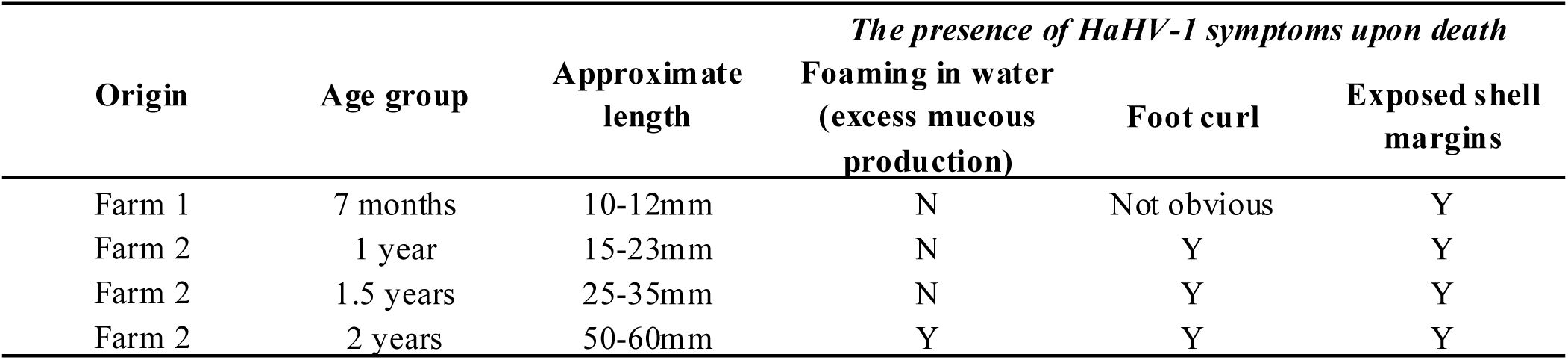
Table key: Y and N indicate the presence and absence of the listed symptom, respectively.

| Origin | Age group | Approximate length | *The presence of HaHV-1 symptoms upon death* |  |  |
| --- | --- | --- | --- | --- | --- |
|  |  |  | Foaming in water<br>(excess mucous production) | Foot curl | Exposed shell margins |
| Farm 1 | 7 months | 10-12mm | N | Not obvious | Y |
| Farm 2 | 1 year | 15-23mm | N | Y | Y |
| Farm 2 | 1.5 years | 25-35mm | N | Y | Y |
| Farm 2 | 2 years | 50-60mm | Y | Y | Y |

### Immune priming against HaHV-1 infection

#### Poly(I:C) priming protects adult abalone from HaHV-1 infection

Currently there is no known treatment for HaHV-1 in abalone, however it has been demonstrated that immune priming with synthetic double stranded RNA (poly(I:C)) can be effective against a related virus, OsHV-1 in oysters^30,28,29^. To investigate if immune priming can protect adult abalone from HaHV-1 infection we initially trailed an intra-muscular injection priming method due to its ease of administration. The survival of 3-year-old hybrid abalone 48hrs post priming with either poly(I:C) or PBS, was measured after HaHV-1 challenge via immersion. As shown in Figure 4A, and although not statistically significant, 50% of poly(I:C) primed abalone survived at the end of the trial as compared to the PBS primed abalone that all died by day 12. To determine if a vascular injection route would improve the anti-viral potential of immune priming against HaHV-1, 3-year-old hybrid abalone were primed with poly(I:C) via injection directly into the pedal sinus (intra-haemolymph poly(I:C)). Survival after challenge with HaHV-1 was measured 5 days post priming and intra-haemolymph poly(I:C) primed abalone maintained a survival rate of 100% throughout the trial (Fig. 4B). This was significantly different when compared to the survival trend of mock-injected abalone which reached ∼80% death and 100% death by day 16 and 24, respectively (Fig. 4B). To understand the longevity of the protection of intra-haemolymph poly(I:C) priming, 3-year-old hybrid abalone were challenged with HaHV-1 via immersion 16-days post priming. As shown in Figure 4C, the survival of poly(I:C) primed abalone by the end of the trial (∼75%) was significantly higher than mock-injected abalone which reached 100% mortality by day 11. The survival of a group of 3-year-old hybrid abalone that was previously intra-haemolymph poly(I:C) primed (126 days prior) and virally challenged via immersion (121 days prior) was also assessed. Interestingly, within this group ∼75% of abalone survived by the end of the trial when compared with the mock primed group, indicating that prior poly(I:C) priming and/or prior exposure to a sub-lethal dose of HaHV-1 protects against subsequent viral challenge (Fig. 4C). To further investigate if poly(I:C) priming is driving an anti-viral response in abalone, a SYBR Green qRT-PCR assay was used to evaluate the transcription of two anti-viral ISGs in abalone haemolymph. A significant foldchange in both viperin and IRF-1 mRNA expression was evident in 3-year-old hybrid abalone that were poly(I:C) primed 2-days prior to sampling as compared to unprimed control abalone (Fig. 4D-E). Additionally, when compared to the control group, the fold change of both viperin and IRF-1 mRNA was trending upwards in abalone that were previously primed (88 days prior) and challenged with HaHV-1 (83 days prior) (Fig. 4D-E).

**Figure 4.**
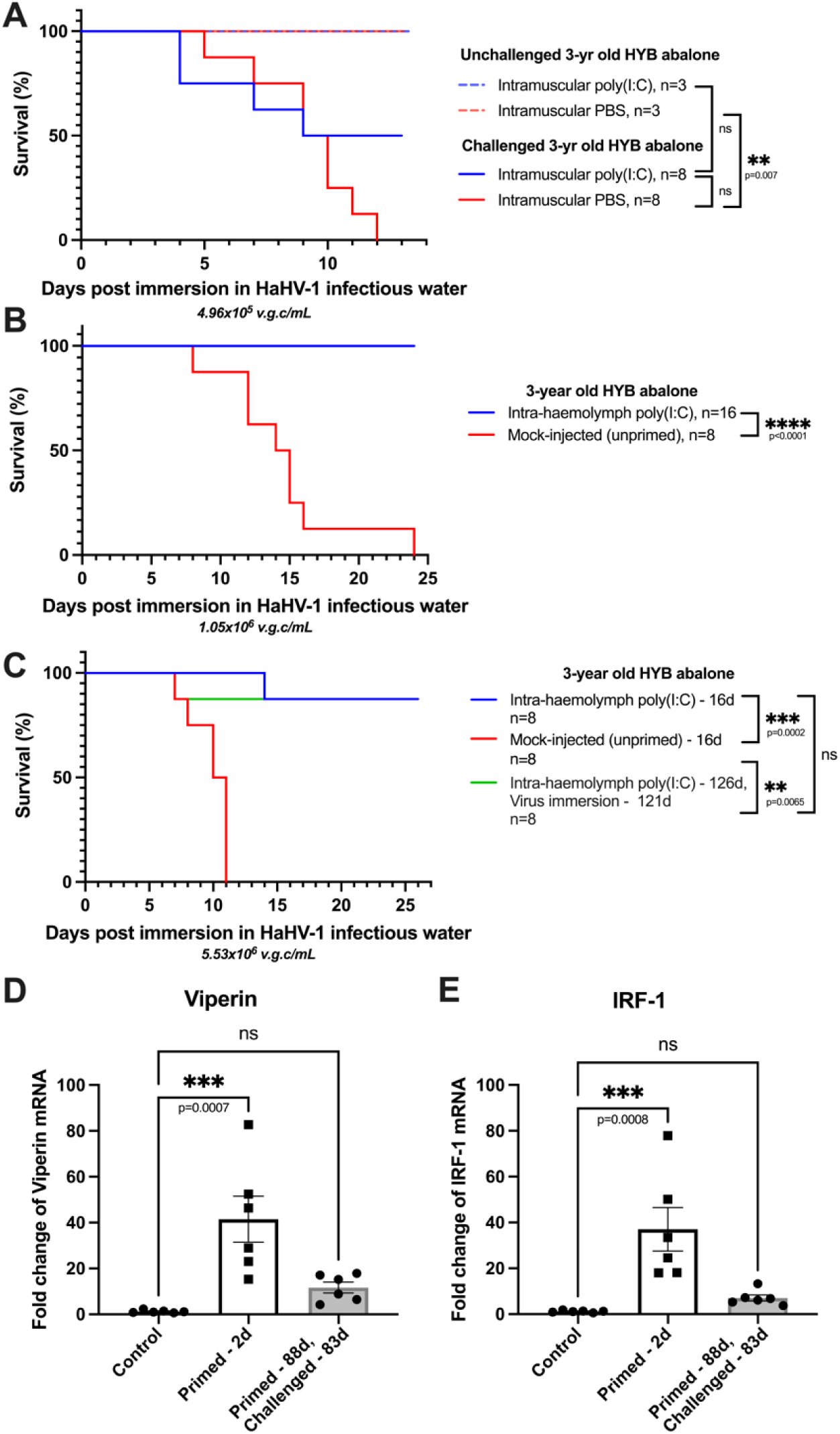
Immune priming protects abalone from HaHV-1 infection. **(a)** Survival of unchallenged (dotted lines) and HaHV-1 challenged (solid lines) 3-year old immune primed (blue lines) and PBS primed (red line) abalone. Abalone were primed with either PBS or poly(I:C) intramuscularly, 2-days prior to infection with ∼4.96 x 10^5^ v.g.c/mL via immersion challenge. Log-rank test, n=3-8. **(b)** Survival of HaHV-1 challenged 3-year old immune primed (blue line) and mock-injected (red line) abalone. Abalone were primed with poly(I:C) or mock-injected via an intra-haemolymph injection, 5 days prior to infection with ∼1.05 x 10^6^ v.g.c/mL via immersion. Log-rank test, n=8-16. **(c)** Survival of HaHV-1 challenged 3-year old immune primed and mock-injected abalone. Abalone were primed with either poly(I:C) (blue line) or mock-injected (red line) via an intra-haemolymph injection prior to infection with ∼5.53 x 10^6^ v.g.c/mL via immersion. An additional group of abalone previously poly(I:C) primed 126 days prior that had survived primary HaHV-1 challenge 121 days prior were rechallenged (green line) with ∼5.53 x 10^6^ v.g.c/mL via immersion. Log-rank test, n=8. **(d-e)** mRNA expression of ISGs, Viperin and IRF-1 in the haemolymph of 3-year old immune primed and un-primed abalone. Abalone haemolymph was sampled 2 days post intra-haemolymph poly(I:C) injection or 88 days post intra-haemolymph poly(I:C) injection in a group of abalone that had previously survived HaHV-1 challenge 83 days prior. Data presented as mean ± SEM, n=6 abalone, ordinary one-way ANOVA, ns= not significant (p>0.05).

#### Abalone primed with recombinantly produced flagellin are not protected from HaHV-1 infection

We next aimed to establish if alternate immune primers known to be anti-viral against other DNA viruses in aquaculture settings, were effective at preventing HaHV-1 infection ^32^. After successful expression, flagellin-A originating from *Vibrio harveyi* was purified from ClearColi™ cells by affinity chromatography (Ni-NTA). As shown in Figure 5A, the molecular weight and purity of the eluted protein was assessed using gel electrophoresis and a single band was detected by SDS-PAGE (41.07 kD). To assess if flagellin-A was protective against HaHV-1, the survival of 2-year-old hybrid abalone was measured 3 days after intra-haemolymph priming with flagellin-A at a high (0.5 mg/mL) and low (0.1 mg/mL) concentration. The trends in survival indicated that flagellin-A priming was not protective against HaHV-1, whereby both primed groups experienced 100% mortality by day 6 post challenge, following a similar trend to the unprimed positive control group and mock-primed group which also displayed 100% mortality by day 7 (Fig. 5B). To understand if flagellin-A could drive an anti-viral response in abalone, the transcription of two anti-viral ISGs was measured in haemolymph from abalone primed 3-days prior with a low, mid and high dose of flagellin-A. Viperin mRNA expression in all flagellin-A primed abalone was not significantly different when compared to unprimed controls (Fig. 5C). Interestingly, IRF-1 mRNA expression was significantly lower in all flagellin-A primed abalone as compared to the unprimed controls (Fig. 5D).

**Figure 5.**
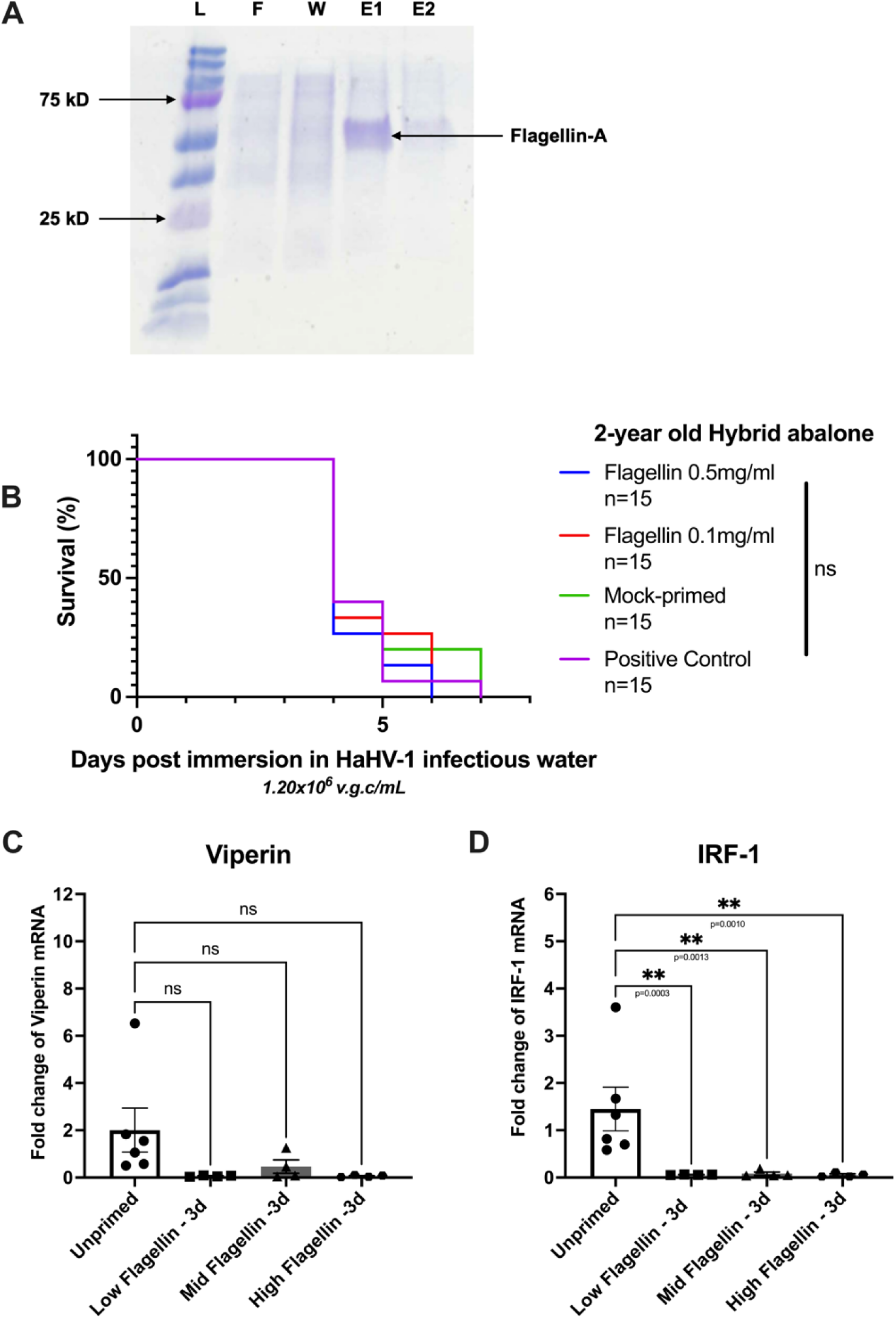
Flagellin-A protein does not protect abalone from HaHV-1 infection. **(a)** Successful Ni-NTA/Ni-IMAC protein purification of *Vibrio harveyi* flagellin-A protein in a Pet28a+ vector in ClearColi™ cells on a Coomassie brilliant blue-stained 12% SDS-PAGE gel. Lane L: Precision Plus Protein Dual Colour Standards (Bio-Rad); Lane F: Flowthrough fraction; Lane W: Wash fraction; Lane E1: First elution fraction; Lane E2: Second elution fraction. **(b)** Survival of HaHV-1 challenged 2-year old flagellin-primed (blue and red lines), mock injected-(green line) and true control (purple line) abalone. Abalone were mock-primed (green line) or flagellin-primed with concentrations of 0.5 mg/mL (blue line) or 0.1 mg/mL (red line) via an intra-haemolymph injection, 3 days prior to infection with ∼1.20 x 10^6^ v.g.c/mL via immersion. Log-rank test, n=15. **(c-d)** mRNA expression of ISGs, Viperin and IRF-1 in the haemolymph of 2-year old flagellin primed- and un-primed abalone. Abalone haemolymph was sampled 3 days post intra-haemolymph-low (10 µ/mL), mid (50 µg/mL) and high flagellin (100 µg/mL) injection. Data presented as mean ± SEM, n=4-6 abalone, ordinary one-way ANOVA, ns= not significant (p>0.05).

### Pedal swabs are a less invasive way to confirm HaHV-1 infection in sick abalone

Confirming HaHV-1 infection in individual abalone during antiviral trials generally requires sampling nerve tissues which necessitates sacrificing the animal, diminishing the ability to longitudinally track animals. To investigate a less invasive method of sampling, HaHV-1 challenged abalone aged 6-months to 1.5-years were swabbed for subsequent viral quantification upon exhibiting symptoms, and their levels compared to nerve samples. All 6-month-old abalone challenged with HaHV-1 showing symptoms of infection had detectable levels of virus ranging from 10^6^-10^11^ v.g.c/mL in swab fluid, in comparison to healthy control abalone, which had undetectable levels of virus (Fig. 6A). We next wanted to assess how resulting viral titres from swabs compare to those of traditional nerve samples of the same animal. Comparable to the above, all 1.5-year-old abalone challenged with HaHV-1 which showed symptoms of infection had detectable levels of virus ranging from 10^6^-10^11^ v.g.c/mL of swab fluid, as compared to healthy control abalone which had undetectable levels of virus (Fig. 6B). Within the nerves however, there was a narrower range of viral titres (10^7^-10^10^ v.g.c/mg) and upon performing correlation analysis there was no correlation (R^2^ = 0.01092) between the viral titre of the HaHV-1 positive swabs and nerves from the same animal (Fig. 6B-C); indicating that pedal swabbing can detect HaHV-1 positivity but is not reliable to determine comparative viral loads between animals.

**Figure 6.**
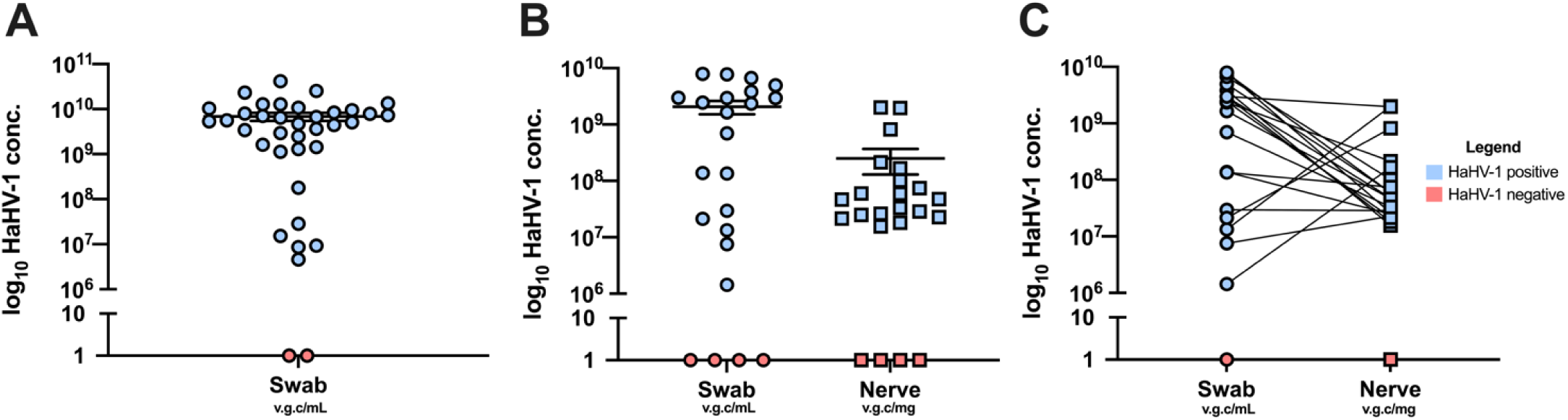
Viral quantification of pedal mucous swab samples in comparison to nerve samples. **(a)** The pedal muscle of 6-month old abalone (n=35) were swabbed upon showing symptoms of HaHV-1 infection after challenge via immersion with ∼1.30 x 10^6^ v.g.c/mL (blue circles). Un-challenged control abalone (n=2) represented by coral coloured circles. Data is represented as mean ± SEM. **(b-c)** The pedal muscle of 1.5-year old abalone (n=19) were swabbed (blue circles) and sacrificed to extract nerves (blue squares) upon showing symptoms of HaHV-1 infection after challenge via immersion with ∼1.44 x 10^6^ v.g.c/mL. Un-challenged control abalone (n=4) are represented by coral coloured circles and squares. Data is represented as mean ± SEM. **(c)** Lines between swab and nerve data points match samples from the same animal. Pearson correlation coefficient analysis, no correlation (R^2^ = 0.01092).

## Discussion

HaHV-1 is a viral pathogen that causes significant mortality events in various key abalone species globally, however further research is required to enhance preparedness for future outbreaks and to identify potential anti-viral preventive measures. This study has revealed several key findings on the susceptibility of Australian hybrid abalone to Haliotid herpesvirus (HaHV-1). Firstly, it was observed that abalone less than one year of age are significantly less susceptible to HaHV-1 infection and exhibit less pronounced clinical signs of infection compared to older abalone. Additionally, immune priming with poly(I:C) in adult abalone provided protection against HaHV-1, whereas priming with Flagellin-A protein did not confer any viral protection. Furthermore, the use of pedal swabs was found to be an effective method for detecting HaHV-1 infection, though it was not reliable for determining comparative viral loads. These findings add critical knowledge and tools that may underpin new preventative strategies against HaHV-1 in aquaculture and emphasise the need for further research on immune priming and age-related susceptibility in abalone.

There is currently a limited understanding of the resistance of juvenile abalone to HaHV-1. Based on published field observations of the 2005 Victorian outbreak, on farm mortalities affected juvenile abalone from 1 year of age to mature broodstock ^7^, however this report did not specify if there was a difference in symptoms or death rate between the affected age groups. Additionally, an Australian study found that wild mature blacklip abalone and blacklip juveniles within the geographical range of AVG along Victoria’s coast had similar survival rates with both groups reaching 100% mortality by approximately day 10 post infection ^21,33^. However, it should be noted here that the “juveniles” in this case were approximately 70 mm in length indicating that these abalone were most likely older than the age we observe the highest resistance to HaHV-1 in our study, which is 7-12 months of age. The resistance of blacklip, greenlip and hybrid abalone to HaHV-1 in this study contrasts with what is known for OsHV-1 infection in oysters ^4,34–36^. OsHV-1 is the only other viral member of the family of molluscan herpesviruses, *Malacoherpesviridae,* but is only distantly related to HaHV-1 ^37–39^. In the economically important pacific oyster (*Crassostrea gigas*), high mortality rates are experienced by oyster spat and juveniles infected with OsHV-1 and its microvariants in both the field and controlled laboratory experiments, which is the reverse of what was observed in abalone in the current study ^4,34,35^. Interestingly however, adult Portuguese oysters (*Crassostrea angulata*) and Blood ark shell clams (*Scapharca broughtonii*) have been shown to be equally or more susceptible to death due to OsHV-1 infection when compared with juvenile animals ^40,41^. This emphasises that susceptibility and mortality trends in one species of mollusc can differ when investigated in not only other families but also other species of the same mollusc. To our knowledge, this is the first time that differential susceptibility based on age has been shown in Australian abalone and highlights the need for controlled studies with a range of species and ages to better understand HaHV-1 mortality dynamics and properly inform the Australian abalone industry for control of future outbreaks.

Immune priming abalone in this study with synthetic nucleic acid prior to viral challenge was protective against HaHV-1 infection and was associated with an anti-viral immune signature when delivered via injection directly into the abalone’s sinus. This result is comparable to trials conducted in the oyster whereby significant protection has been evidenced against OsHV-1 challenge in both laboratory and wild settings upon direct injection of poly(I:C) into the adductor muscle ^28–30,42^. The adductor muscle of a bivalve organism is highly vascular and therefore a direct route into the circulation which we were able to mirror in the gastropod abalone through priming directly into the pedal sinus ^43,44^. Additionally, within this current study, poly(I:C) priming abalone 16 days prior to primary HaHV-1 challenge and 126 days prior to secondary HaHV-1 challenge, significantly protected abalone from infection when compared to mock handled controls. This outcome suggests that priming abalone with poly(I:C) may afford longitudinal protection, similarly to studies conducted in the oyster, where protection against OsHV-1 was present for a total of five months post priming, in both a laboratory and field based environment ^28^. In the group that was primed 126 days prior to secondary HaHV-1 challenge, it is unclear if protection was afforded solely by poly(I:C) priming, the initial subclinical HaHV-1 challenge or a combination of the two as exposure to sublethal doses of viruses can act as an immune priming event ^25^. Regardless, the significant upregulation of transcripts relating to abalone ISGs, viperin and IRF-1, 88 days post prime, compared to an untreated control support the longevity of the afforded protection. Similar studies previously conducted in the oyster have also demonstrated the upregulation of such immune related genes and pathways highlighting the ability of this priming agent to initiate and prolong an anti-viral state ^29,30,45,46^. To understand if the anti-viral protection afforded by poly(I:C) could be achieved by an alternate immunostimulant, flagellin-A was also trialled as an immune primer against HaHV-1. Flagellin is a highly conserved bacterial protein that is often utilised as a broad-spectrum adjuvant in vaccines due to its pro inflammatory nature and has recently been found to facilitate an anti-viral response against white spot syndrome virus (WSSV) in shrimp ^32,47^. In this current study, priming abalone with flagellin-A did not have an anti-viral effect and was not capable of providing protection against HaHV-1 infection which may indicate that the abalone’s innate immune response to flagellin is different to that of other aquatic species ^48^. Consequently, poly(I:C) priming appears to provide abalone with long-term protection against HaHV-1, however, in depth molecular analysis of the basis of immune priming in these organisms is required to draw meaningful conclusions on the immune modulation of this species upon treatment with poly(I:C) to inform future immune priming studies.

Detecting HaHV-1 infection early may facilitate outbreak control, however current methods require sampling nervous tissue ^15,18^. This study showed that swabbing suspected HaHV-1 positive abalone offers a less invasive and more rapid method to confirm infection, even in abalone as young as six months old. The ability to more rapidly detect the virus without invasive sampling methods that kill the animal, will also allow for longitudinal tracking of the infection in the same specimen to further our understanding of the pathogenesis of this virus. The use of alternative pathogen detection methods such as loop-mediated isothermal amplification (LAMP) assays has steadily increased due to their ability to provide on-site detection in an accurate, rapid, and cost effective manner ^49^. An efficient and specific LAMP assay has been developed for HaHV-1 but has only been optimised for pathogen detection with DNA extracted from tissue ^50^. To accommodate the on-site application of LAMP, a farm deployable method of DNA extraction from a non-invasive sampling site is ideal. A cost efficient and rapid DNA extraction method from non-invasive mucosal swabs has recently been optimised in crucian carp (*Carassius auratus*) ^51^. This swabbing and DNA extraction technique was effective at producing sufficient DNA for subsequent PCR amplification and the identification of positive and negative *C. auratus* herpesvirus (CaHV) infections ^51^. Furthermore, a portable DNA extraction technique optimised for use in LAMP assays has also been developed for WSSV in shrimp, highlighting the potential of optimising a similar viral identification strategy for HaHV-1 in farmed abalone ^52^. The ability to quickly and accurately identify active HaHV-1 infections on-farm without specialised equipment or expertise, is critical to inform the effective control of viral spread both throughout the farm and to coastal wild abalone, given that farm outflow water is returned to adjacent water bodies.

Abalone herpesvirus is a significant virus of concern for both abalone farmers and the wild sector, with recurrent outbreaks over the last few decades in southern regions of Australia. There are currently no strategies to safeguard farmed abalone against this virus, and further work to better understand the pathogenesis of the virus and its control may assist in the development of novel strategies to protect farmed abalone. This study describes that there is an effect of abalone age on HaHV-1 susceptibility and demonstrates for the first time that immune priming abalone with poly(I:C) prior to challenge, protects them from future infection. This research coupled with the new knowledge that pedal swabs with subsequent nucleic acid testing give an accurate diagnosis of HaHV-1 will support future endeavours to develop novel strategies to protect abalone against HaHV-1 infection.

## Acknowledgements

This work was funded by ‘2021-085 Optimizing an immune priming approach to combat HaHV-1 in abalone’, supported by funding from the FRDC on behalf of the Australian Government to KJH and TB. JA and DA are supported by a La Trobe University industry PhD scholarship in conjunction with the abalone growers association of Australia. We also wish to acknowledge the staff at the abalone farms involved in this project, the technical staff of the School of Agriculture, Biomedicine and the Environment and the members of the Helbig Lab.

## Author Contributions

JR Agius: Writing – original draft, Investigation, Data Curation, Methodology, Writing – review and editing. D Ackerly: Writing – original draft, Investigation, Data Curation, Methodology. AC Watson: Investigation, Writing review and editing. ML Smith: Investigation, Writing review and editing. T Beddoe: Conceptualization, Funding acquisition, Supervision, writing – review and editing. KJ Helbig: Conceptualization, Funding acquisition, Supervision, Writing – original draft, Writing – review and editing.

## Data Availability Statement

All data is available upon request.

## Methods

### Experimental animals and general maintenance

All abalone included in this study were kindly gifted by collaborating Victorian abalone farms located in Avalon (Farm 1), Indented Head, Port Fairy (Farm 2) or Narrawong (Farm 3). Experiments were performed with either greenlip (*Haliotis laevigata)* or hybrid abalone *(H. rubra X H. laevigata)* aged 7-months to 3-years of age. Abalone were acclimatised in the laboratory at least 1-2 weeks prior to priming and challenge experiments. Abalone were co-housed in groups at 17±1 °C in an appropriate volume of aerated natural seawater (sourced from Mount Martha) which was calculated based on previous optimisation and the average mass of the abalone (∼32.10 mL seawater/g of abalone ± 25%). Unless otherwise stated, 100% water changes were conducted at least once a week to maintain water quality. In general, abalone were not fed throughout the duration of the viral trial, however feeding did occur during the acclimatisation period.

### Upscaled and purified HaHV-1 viral homogenate

All viral stocks utilised to produce HaHV-1 infectious water were upscaled from archival snap-frozen abalone tissue sourced from the 2005 Victorian outbreak (HaHV-1 VIC-1), and acquired from the Department of Jobs, Precincts and Regions (DJPR). Once thawed, extracted nerve tissue was homogenised on ice in 1mL of minimal essential medium (MEM, Thermo Fisher, Cat: 11095098). The homogenate was transferred into a 15mL falcon tube and centrifuged at 1500 x g for 20 min at 4 °C, in a Velocity 18R Versatile Centrifuge. The supernatant containing the virus was removed and filtered (0.45 µM) into a fresh tube. FBS and glycerol were added to the viral stock to final concentrations of 10% (v/v) and 1% (v/v) respectively which was then stored in aliquots at −80 °C. To confirm the absence of culturable bacteria, an aliquot of the viral stock was plated on Luria Broth agar which was incubated at 18 °C for one week. For subsequent upscaling, 200µL of this HaHV-1 stock was injected intramuscularly (25-G needle fitted to 1mL syringe) into 3 healthy abalone which were then housed together at 18 °C in seawater aerated with an air stone. Once abalone were moribund and exhibiting HaHV-1 symptoms, they were dissected as demonstrated in the AQUAVETPLAN ^18^ to isolate nervous tissue which was then pooled, prepared and stored as above to produce a P1 infectious HaHV-1 homogenate. To titre the viral stock, DNA was extracted using a ReliaPrep gDNA Tissue Miniprep System (Promega, Cat: A2051) following the manufacturer’s instructions for a buccal swab. The samples were eluted in 50 µL nuclease free water and stored at 4 °C short-term, until qPCR analysis as described below. All following HaHV-1 viral homogenates were generally prepared as described above with some minor adjustments in number of abalone used, infection route (immersion instead of injection) and media (DMEM instead of MEM).

### Preparation of HaHV-1 infectious water and co-housed immersion challenge model

HaHV-1 infectious water and the immersion challenge was adapted from ^53^ and is described in Figure 1. Prior to each infection trial, abalone (hybrid abalone of approximately 2-3 years old) were intramuscularly injected (25-G needle fitted to 1mL syringe) with ∼5.56 x 10^7^ v.g.c/mL (diluted with DMEM) of HaHV-1 viral homogenate into each side of the foot (100 µl viral inoculum per side). Abalone were then co-housed in fresh seawater aerated with an air stone, with a water change 24hrs following injection until they presented as moribund to allow maximum viral shedding into the water, typically 4 days. Abalone were removed from the water when they became moribund, and a sample of the infectious water was collected to determine the relative viral titre via DNA extraction and qPCR analysis as described below. In the required aquaria, excess water was removed and replaced with titred infectious water to achieve a desired infection dose. To simulate the natural pressures of HaHV-1 infection outside of the lab, abalone were co-housed during and after the immersion process. A 100% water change was conducted on all aquaria 24hrs post immersion and 48hrs thereafter. Abalone health was monitored daily and presentations of clinical signs of HaHV-1 infection such as the loss of adherence to the substrate, foot curl, prolapse of the mouth and the presence of excessive foaming was recorded. Once no longer able to adhere to the substrate and turn themselves upright, abalone were recorded as dead and removed from the aquaria. All co-housed groups challenged with HaHV-1 via immersion had a matched negative co-housed control group that did not receive virus.

### Poly(I:C) priming solution

High molecular weight poly(I:C) (InvivoGen, Cat: tlrl-pic-5) was made up in endotoxin free water (1 mg/mL) as per the manufacturer’s instructions and stored at −20⁰C until use. Abalone were placed shell down on ice lined with paper towel (∼2 min), to minimise stress and were then injected with thawed poly(I:C) at a final concentration of 100 µg per abalone with 25-gauge needles fit to 1 mL syringes.

### Flagellin-A priming solution

#### Gene cloning and Plasmid construction of Vibrio harveyi flagellin A gene

The flagellin A sequence form *Vibrio harveyi* strain (1DA3) (NCBI accession no. EEz87745.1) was codon-harmonised and commercially synthesised and cloned into pET28a(+) vector (Novagen, Sigma-Aldrich, Cat: 69864-3). Restriction enzymes, Ndel and XhoI were used to cleave the flagellin-A sequence and the pET28a(+) vector between N and C-terminal polyhistidine tag (His tag) with a stop codon added to the sequence upstream of the C-terminal his tag.

#### Transformation and expression of recombinant flagellin A

The recombinant flagellin-A pET28a(+) construct was initially transformed into chemically competent T7 Express *lysY/I^q^* Competent *E. coli* (High Efficiency) cells (New England Biolabs, Cat: C3013I) according to standard heat shock *E. coli* transformation protocols ^54^. Successful transformants were identified by growth on Luria Bertani (LB) agar plates (1% (*w/v*) Tryptone, 0.5% (*w/v*) Yeast Extract, 1% (*w/v*) NaCl, 1.5% agar, pH [7.4]) containing 50 µg/uL Kanamycin.

Successful transformants were inoculated in 10 mL LB broth containing 50 µg/mL of Kanamycin and incubated at 37 °C for 16 hrs at 225 rpm. The flagellin-A protein was expressed in 400 mL of LB media containing 1:100 dilution of the 10 mL seed culture within a 2 L baffled flask. Cultures were incubated at 37 °C with shaking at 200 rpm until the Optical Density at 600 (OD600) of 0.4-0.6 was reached and protein expression was induced by 1 mM Isopropyl β-D-1-thiogalactopyranoside (IPTG, GoldBio, Cat: I2481C). Proteins were incubated for an additional 3 hrs to maximise protein expression and harvested by centrifugation at 7000 x g in 4 °C for 20 min at 4 °C. Cell pellets were lysed in 30 mL *E.coli* lysis buffer (20 mM Tris-HCl, pH [8.0], 300 mM NaCl, 5 mM Imidazole with 0.1 mg/mL lysozyme) and disrupted by sonication with 4 cycles of 35 amps for 30 seconds alternated by 30 seconds of intermittent cooling on ice for a total of 4 min (VCX 130, VibraCell, Sonics). Cellular debris was removed by centrifugation at 18,000 rpm for 20 min at 4 °C with bacterial cells removed from cell lysate by passing culture through 0.45 µM filter and stored at 4 °C.

#### Protein purification and Separation via SDS-PAGE and Western Blot Analysis

The cell lysate obtained after sonication and filtration was purified using nickel-nitrile acid (Ni-NTA) affinity chromatography. Briefly 2 mL of concentrated nickel agarose resin (QIAGEN, Hilden, Germany, Cat:30210) was added to a gravity column and equilibrated with 20 column volumes (CV) of *E. coli* lysis buffer at a flow rate of 1 mL/min under gravity pressure. The cell lysate was added to the column at the same flow rate of 1 mL/min with 1 mL aliquot flow through solutions collected each purification step for SDS-PAGE gel analysis. Then the column was washed to remove unbound proteins with a wash buffer (50 mM NaH2PO4, 300 mM NaCl, 20 mM Imidazole, pH [7.4]. Bound proteins were eluted in a starter elution buffer (50 mM NaH2PO4, 300 mM NaCl, 300 mM Imidazole, pH [7.4] then eluted in a final elution buffer with 500 mM Imidazole. Elution’s were combined and buffer exchanged into 1 x PBS (120 mM NaCl, 5 mM NaH2PO4 H2O, 16 mM Na2HPO4, pH [7.4] using Amicon®Ultra-15 centrifugal filters units (Merck, Kenilworth, IL, USA, Cat: UFC900308) with a 3 kDa cut-off. Protein concentrations were determined using the Pierce BCA Protein Assay Kit (Thermo Fisher Scientific) according to manufacturer’s instructions.

Purified samples were analysed by Sodium dodecyl sulphate–polyacrylamide gel electrophoresis (SDS-PAGE) separation using a 12% TGX Stain-Free Fast Cast Acrylamide kit as per manufacturer’s instructions (Bio-Rad, California, USA, Cat: 1610185) and stained with 0.1% (w/v) Coomassie brilliant blue dye. All samples were mixed with 2X Laemmli sample buffer and boiled for 10 min at 95 °C prior to centrifugation at 13,000 rpm for 5 min. Gels were run at 200V for 35 min in 1X SDS-Running buffer consisting of 3.0 % (w/v) Tris-HCl, 14.4 % (w/v) Glycine, 1% (w/v) SDS, pH [8.3] in Mini-Protean Tetra cells (Bio-Rad, Cat: 1658005EDU) on a PowerPac™ Basic Power Supply (Bio-Rad, Cat: 1645050) and after staining were imaged on a CanoScan (Cannon, Cat: N67OU). SDS-PAGE gels were replicated with one stained with Coomassie and the other used for Western blot analysis. For Western blot analysis, proteins were electro-transferred onto a PVDF membrane for 7 min as per manufacturer’s instructions utilising the Trans-Blot ® Turbo™ System (Bio-Rad, Cat: 1704150). Membranes were blocked with a blocking buffer (5% (w/v) skim milk in (0.05% (w/v) in TBS-T: 50 mM Tris-HCl, pH [7.6], and incubated for 1 hr at ambient temperature with orbital shaking. Membranes were then washed in TBS-T prior to incubation with mouse anti-His tag (R & D systems, Cat: MAB050) antibodies diluted to a 1:10,000 in blocking buffer for 1 hr at ambient temperature for 1 hr with orbital shaking. Membranes were washed in TBS-T prior to final development with the Clarity Western ECL™ substrate as per manufacturers instruction (Bio-Rad, Cat: 1705061) and visualised on the C-DiGit® Blot Scanner (LI-COR Biosciences, Lincoln, NE, USA).

#### Transformation and expression of recombinant flagellin A in LPS free cell line

Purified flagellin-A pET28a(+) plasmid was transformed into electrocompetent Lipopolysaccharide (LPS) free cell line, ClearColi™ BL21 (DE3) (Lucigen, Cat: 60810-1) according to manufacturer’s recommendations ^55^. Briefly, 1 µL of flagellin-A pET28a(+) plasmid was combined with 25 µL of ClearColi™ BL21 (DE3) cells into a 0.1 cm gap electroporation cuvette (Biorad, Cat: 1652083) and given a single pulse optimised for *E.coli* cells at 1.8 kV, a capacitance of 25 µF and 200 ohms using the Gene Pulser Xcell Electroporation Systems (Biorad, Cat: 1652660). Transformed cells were recovered in 975µL of expression recovery medium for 1 hr at 37 °C with shaking at 250 rpm then plated on LB agar with 50 µg/µl of Kanamycin and incubated at 37 °C for 40 hrs to obtain single colonies.

Single colonies were inoculated in 10 mL LB broth containing 50 µg/mL of Kanamycin and 0.5% (w/v) glucose (Sigma-Aldrich, Cat: G8270) to minimise expression and incubated at 37 °C for 16 hrs at 225 rpm. The flagellin-A protein was expressed in 400 mL of LB media containing a 1:100 dilution of the 10 mL seed culture, incubated at 37 °C with shaking at 200 rpm until an OD600 of 0.6-0.8 was reached. At this OD600, protein expression was induced by 1 mM Isopropyl β-D-1-thiogalactopyranoside (IPTG, GoldBio, Cat: I2481C) and incubated for a further 4 hrs to maximise protein expression. Cells were harvested by centrifugation at 5000 x g in 4 °C for 10 min at 4 °C. Proteins were extracted and purified as previously described and separated by SDS-PAGE analysis

### Priming route

#### Intra-muscular priming

Preparations for intra-muscular poly(I:C) priming were performed as described above with minor adjustments to make the final volume in each syringe 200 μL (500 μg/mL poly(I:C) in sterile 2 x phosphate-buffered saline (PBS)). Abalone were intra-muscularly injected with 100 μL of poly(I:C) into each side of the foot. Un-primed controls were handled in the same way but were intra-muscularly injected with a total of 200 μL 1 x PBS. Once injected, abalone were returned to their respective aquaria and monitored until 100% adherence to the surface was observed in preparation for subsequent immersion challenge in either titred HaHV-1 infectious water or virus free natural seawater.

#### Intra-haemolymph priming

Preparations for intra-haemolymph poly(I:C) priming were performed as described above with minor adjustments to make the final volume 100 μL in each syringe (1 mg/mL). Abalone were injected with a pre-filled poly(I:C) syringe directly into the abalone’s pedal sinus located in the midline of the foot, approximately 2 cm from the head. To confirm the syringe was correctly inserted into the sinus, the syringe was drawn until haemolymph was extracted into the barrel. Without removing the syringe, the poly(I:C) and haemolymph mixture was slowly returned to the animal and light pressure was placed on the site of injection upon removal of the needle. Un-primed controls were mock-injected in the same location by drawing up haemolymph into an empty syringe and returning the contents to the animal. Once injected, abalone were returned to their respective aquaria and monitored until 100% adherence to the surface was observed in preparation for subsequent immersion challenge in either titred HaHV-1 infectious water or virus free natural seawater.

### Sampling of abalone and water for viral detection and quantification

DNA extractions were conducted with a ReliaPrep gDNA Tissue Miniprep System (Promega, Cat: A2051) following the manufacturer’s instructions with minor adjustments dependent on the sample type. All samples were eluted in 50 µL nuclease free water and stored at 4 °C short-term, until qPCR analysis. Prior to extractions, all water samples were centrifuged at maximum speed for 5 min in biocontainment buckets in a Velocity 18R Versatile Centrifuge. DNA extraction was then performed on 100 μL of the supernatant following the manufacturer’s instructions for a buccal swab. For nerve DNA extractions, a small square portion of nervous tissue was dissected just below the abalone’s head when the pedal nerve cords were visible as 2 distinct dots. The maximum amount of muscle was removed from the portion which was then weighed and extracted following the manufacturer’s standard protocol for animal tissue. For swab samples, the surface of the pedal muscle was swabbed using one sterile cotton swab applicator per abalone with its handle snapped off to fit in a closed 1.5 mL Eppendorf tube. Abalone were initially swabbed three times around the circumference of the lip, with the swab being rotated during each pass. The swab was then rotated up and down the midline of the abalone’s foot 3 times and then rotated around the abalones mouth for 2 rotations. The swab was then placed in a 1.5 µL tube pre-filled with 400 µL of PBS, stick end pointing up. The manufacturer’s DNA extraction protocol for buccal swabs was then followed from step 2.

### Viral HaHV-1 quantification via quantitative PCR

The relative titre of HaHV-1 DNA was quantified using qPCR primers targeting the open-reading frame (ORF) 66 (Forward - 5’ TCCCGGACACCAGTAAGAAC & Reverse - 5’ CAAGGCTGCTATGCGTATGA) of the Victorian isolate of HaHV-1 as it is outlined to be effective in HaHV-1 disease investigations in both the AQUAVETPLAN and the World Organisation for Animal Health (OIE) guidelines ^15,18^. Each 20 μL qPCR reaction contained: 0.3 μL of the forward and reverse primers (20 pmol/mL), 5 μL of SensiFAST SYBR No-ROX Kit (Bioline, Cat: BIO 98020), 9.4 μL of nuclease free H_2_O and 5 μL of extracted DNA. The amplicon length of ORF66 was 145 base pairs (bp). Reaction cycling conditions consisted of denaturation at 95 ⁰C for 59 sec, followed by 45 cycles of 95 ⁰C for 3 sec and 60 ⁰C for 30 sec. For analysis of the melt curve, a step of 65 ⁰C for 5 sec followed by incremental increases of 0.5 ⁰C every 5 sec, until a temperature of 95 ⁰C was reached was utilised. Analysis of PCR products was performed on a CFX Connect Real-Time PCR Detection System (BioRad). Relative viral titres were determined from the Ct values produced through the preparation of a standard curve.

ORF66 was PCR amplified using the extracted viral DNA from the upscaled stock with a GoTaq G2 Flexi (Promega, Cat: M7801) polymerase using primers (Forward: 5’ ATGGGAATTCTCCCGGACACCAGTAAGAAC & Reverse: 5’ GCTGTCTAGAC AAGGCTGCTATGCGTATGA) as per manufacturer’s instructions. Reaction cycling conditions consisted of initial denaturation 95 ⁰C for 2 min, followed by 34 cycles of 95 ⁰C for 30 sec, 59 ⁰C for 30 sec and 72 ⁰C for 30 sec followed by a final extension of 72 ⁰C for 5 min. PCR products were run on a 2 % agarose gel and then a gel clean-up was performed with an Isolate II PCR and Gel Kit (Bioline, Cat: BIO-52059). The resulting PCR products and plasmid backbone (pcDNA3) were then digested with EcoRI-HF and XbaI for 3 hours at 37 ⁰C (NEB, Cat: R3101S; Cat: R0145S) to be run on a gel and cleaned up as described above. The resulting products were then ligated together with T4 DNA ligase (NEB, Cat: M0202S) and transformed into competent DH5α cells. Five single colonies were selected from each transformation and mini-DNA extractions were performed utilising in-lab kits. Successful cloning of ORF66 into a pcDNA3 vector was confirmed via Sanga sequencing. Plasmids were quantified utilising the A260 measurement determined by a NanoPhotometer Classic (Implen). Once quantified, the copy number of the 5553 bp construct per unit mass for ORF66 was calculated using the formula: number of copies = (DNA concentration x Avagadro’s number)/(bp length x 1 x 10^9^ x 650) ^56^. A standard curve was generated by amplifying a series of 10-fold dilutions ranging from 10^8^ to 10^2^ ORF 66 copies/μL in duplicate via qPCR. To visualise the standard curve, the copy numbers were plotted against the generated cycle threshold (Ct) value. Copy numbers (henceforth referred to as viral gene copies or v.g.c) of samples could therefore be quantified by comparing their Ct values with the standard curve thus obtaining the relative quantity of HaHV-1 viral gene copies present.

### Sampling of abalone haemolymph

Abalone were placed shell down for approximately 2 min on ice lined with paper towel. Using one 25-gauge needle fitted to 1 mL syringe, haemolymph was drawn from the pedal sinus. The needle of the syringe was removed, and the contents then expressed into a clean Eppendorf tube on ice for immediate processing. A new needle and syringe were utilised for each individual animal requiring haemolymph sampling.

#### RNA extraction

For quantitative mRNA expression studies, total cellular RNA was extracted from haemocytes utilising TRIsure Reagent (Bioline, Cat: BIO-38033) to lyse the cells. Each Eppendorf tube containing 150 μL of freshly extracted haemolymph was treated with 850 μL TRIsure and incubated at room temperature for 5 min. 200 μL of chloroform was then added to each sample, mixed thoroughly, and incubated at room temperature for 5 min prior to being centrifuged at 12,000 x g at 4 °C for 30 min. The top aqueous layer was transferred into a fresh Eppendorf tube without disturbing the other layers. Total RNA was precipitated by first adding 1 μL of glycogen (20 mg/mL, Roche, Cat: 10901393001), flicking to mix and subsequently adding 500 μL of cold isopropanol, tilting to mix. Samples were incubated for ∼10 min and centrifuged at 15,000 x g at 4 °C for 30 min to pellet the RNA. The supernatant was removed, and the pellet was washed once with 1 mL of 75% (v/v) ethanol in RNase-free dH_2_O and centrifuged at 7,000 x g at 4 °C for 5 min. The supernatant was removed, and the pellet was airdried (∼10 min) by inverting the tube onto paper towel. Once dry, the pellet was dissolved in 12 μL nuclease free water and stored at −80 °C until further processing.

#### Real-time PCR

First-strand cDNA was synthesised from the total RNA using a Tetro cDNA Synthesis Kit (Bioline, Cat; BIO-65043). The reaction was performed in a pre-chilled RNase-free 0.2 mL PCR tube using the whole volume (12 μL) of previously extracted RNA. Previous experiments have indicated a low yield of RNA in haemolymph, so the purity and quantity of extracted RNA was not assessed prior to synthesising cDNA to maximise the available RNA for cDNA synthesis. The cDNA synthesis reaction mix used with 12 μL total RNA included 1 μL random hexamer primer, 4 μL 5x RT Buffer, 1 μL RiboSafe RNase inhibitor and 1 μL Tetro Reverse Transcriptase (200 U/μL). cDNA was synthesised by incubating the specified amount of total RNA added with cDNA synthesis master mix. First, the synthesis samples were incubated for 10 min at 25 °C followed by 45 °C for 30 min. Then the reactions were terminated by incubating the reaction mix at 85 °C for 5 min followed by chilling on ice. Finally, cDNA samples were diluted to a final volume of 80 μL with DEPC treated H_2_O and stored in −20 °C freezer for long term storage.

qPCR analysis was conducted on the resulting synthesised cDNA to quantify the change in mRNA expression of targeted genes (IRF-1 and viperin) in comparison to the abalone housekeeping gene (RL-7). Each 20 μL qPCR reaction contained: 0.3 μL of each primer (20 pmol/mL), 5 μL of SensiFAST SYBR No-ROX Kit (Bioline, Cat: BIO 98020), 9.4 μL of nuclease free H_2_O and 5 μL cDNA. Reaction cycling conditions consisted of denaturation at 95 ⁰C for 2 min, followed by 39 cycles of 95 ⁰C for 5 sec and 60 ⁰C for 20 sec. For analysis of the melt curve, a step of 65 ⁰C for 5 sec followed by incremental increases of 0.5 ⁰C every 5 sec, until a temperature of 95 ⁰C was reached was utilised. The analysis of PCR products was performed on a CFX Connect Real-Time PCR Detection System (BioRad). Previously published primers were utilised in this analysis and are outlined in Table 2.

### Standard statistical analysis

All statistical analyses presented and plotted using GraphPad Prism version 10.4.1 for Mac (GraphPad Software, San Diego, California USA, www.graphpad.com) and considered significant when the p-value was less than 0.05. Student’s t tests were used for statistical analysis between 2 groups and statistical data from more than 2 groups was analysed with an ordinary one-way ANOVA. Log-rank tests were used for statistical analysis between Kaplan-Meier survival curves.

